# Not all that is gold glitters: PV-IRES-Cre mouse line shows low efficiency of labeling of parvalbumin interneurons in the perirhinal cortex

**DOI:** 10.1101/2021.09.23.461505

**Authors:** Maximiliano José Nigro, Hinako Kirikae, Kasper Kjelsberg, Rajeevkumar Raveendran Nair, Menno P. Witter

## Abstract

The wide diversity of cortical inhibitory neuron types populating the cortex allows the assembly of diverse microcircuits and endows these circuits with different computational properties. Thus, characterizing neuronal diversity is fundamental to describe the building blocks of cortical microcircuits and probe their function. To this purpose, the mouse has emerged as a powerful tool to genetically label and manipulate specific inhibitory cell-types in the mammalian brain. Among these cell-types, the parvalbumin-expressing interneuron type (PV-INs) is perhaps the most characterized. Several mouse lines have been generated to target PV-INs. Among these mouse lines, the PV-IRES-Cre lines is the most widely used and demonstrated a high specificity and efficiency in targeting PV-INs in different cortical areas. However, a characterization of the performance across cortical regions is still missing. Here we show that the PV-IRES-Cre mouse line labels only a fraction of parvalbumin immunoreactive neurons in perirhinal cortex and other association areas. Our results point to a yet uncharacterized diversity within the PV-INs and emphasize the need to characterize these tools in specific cortical areas.

## Introduction

The cerebral cortex is populated by a wide diversity of glutamatergic and GABAergic neurons (Harris and Shepherd, 2015). GABAergic neurons represent 10-15% of the neuronal population, but their diversity allows them to participate in different microcircuits and shape information flow in the cortex (Kepecs and Fishell, 2014). The study of neuronal diversity was propelled by the discovery of molecular markers that are specifically expressed by neuronal classes. GABAergic neurons can be broadly divided in three major classes according to the non-overlapping expression of parvalbumin (PV), somatostatin (SST) and 5-hydroxytryptamine 3a receptor (5HT3aR) (Tremblay et al., 2016). The discovery of molecular markers has been exploited to produce transgenic mouse lines expressing fluorophores or recombinases to label or manipulate specific neuronal classes (Madisen et al., 2010; Taniguchi et al., 2011). Parvalbumin-expressing GABAergic neurons (PV-INs) are the major group of inhibitory neurons in the mammalian cortex, representing about 40% of the GABAergic population (Tremblay et al., 2016). PV-INs provide powerful perisomatic inhibition to excitatory neurons and are involved in feedforward and feedback circuits (Hu et al., 2014; Tremblay et al., 2016). PV-IN encompass two major morphological cell-types: fast-spiking basket cells that target the soma of postsynaptic neurons, and chandelier cells, targeting the axon initial segment. The development of Knock-in mice expressing Cre or FlpO recombinase under the control of the PV promoter (PV-IRES-Cre) has provided researchers with a powerful tool to explore the functional role of PV-INs in the cortical circuit (Hippenmeyer et al., 2005). The PV-IRES-Cre mouse line has been shown to efficiently and specifically target PV-INs in the visual cortex (Atallah et al., 2012), and in the medial entorhinal cortex (Martinez et al., 2017). Recently the PV-IRES-Cre mouse line has been used to quantify the density of PV-INs across the whole cortex (Whissell et al., 2015). These studies highlighted the low density of PV-INs in associative cortices, including the perirhinal cortex. However, a thorough characterization of the specificity and efficiency of the PV-IRES-Cre line in perirhinal cortex is still missing. The perirhinal cortex is part of the parahippocampal network and represents a gateway of sensory information entering the lateral entorhinal cortex (LEC), and the hippocampus (Witter et al., 2000). The perirhinal cortex has been shown to provide an inhibitory control on the information entering the LEC (de Curtis and Paré, 2004). The mechanism underlying this inhibitory control has been hypothesized to be mediated by PV-INs in the deep layers of PER through feedforward inhibition (Willems et al., 2018). This proposed role of PV-INs in gating information flow to the LEC seems to contrast with the low density of labeled cells in PER of the PV-IRES-Cre mouse line. In the current study we quantified the specificity and efficiency of the PV-IRES-Cre mouse line in PER. We report that the PV-IRES-Cre targets only a fraction of PV immunoreactive neurons in perirhinal cortex and other association areas. We suggest that these results point to a yet uncharacterized diversity within the PV-INs.

## Methods

### Animal models

The research described here was performed on commercially available transgenic mice. We used the following mouse lines: PV-IRES-Cre (B6;129P2-Pvalb^tm1(cre)Arbr^/J; stock 008069, The Jackson Laboratory), Ai9 (Gt(ROSA)26SorRCL-tdT; stock 007909, The Jackson Laboratory), R26R-EYFP (129X1-Gt(ROSA)26Sor^tm1(EYFP)Cos^/J (stock 006148, The Jackson Laboratory). The PV-IRES-Cre and Ai9 mice were bred as homozygotes, whereas the R26R-EYFP mice were bred as heterozygotes. To express fluorescent proteins in PV-INs we crossed PV-IRES-Cre mice with either Ai9 or R26R-EYFP mice. Mice were group housed, with water and food ad libitum and a reverse dark/light cycle of 12h/12h. We used adult mice (>2 months) of both sexes in the current study.

### AAV2/1-FLEX-mCherry purification

First, pAAV-CMV-βglobin-intron-FLEX-MCS-WPRE was created by cloning a FLEX cassette with multicloning site into Cla1 and HindIII sites in pAAV-CMV-βglobin-intron-MCS-WPRE (Agilent). Sequence of the FLEX cassette was obtained from (Atasoy et al., 2008). Subsequently, mCherry sequence was synthesized and cloned in the inverted orientation into EcoR1 and BamH1 sites in pAAV-CMV-βglobin-intron-FLEX-MCS-WPRE to create pAAV CMV-βglobin-intron-FLEX-mCherry-WPRE. The positive clones were confirmed by restriction digestion analyses and subsequently by DNA sequencing. Endotoxin free plasmid maxipreps (Qiagen) were used for AAV preparations. Approximately 7× 10^6^ AAV 293 cells (Agilent) were seeded in DMEM containing 10% fetal bovine serum (ThermoFisher) and penicillin/streptomycin antibiotics into 150 mm cell culture plates. Next day, calcium chloride mediated cotransfection was done with 22.5 μg pAAV-containing the transgene, 22.5 μg pHelper, 11.3 μg pRC, 11.3 μg pXR1 (NGVB, IU, USA) capsid plasmids. The medium was replaced by fresh 10% FBS containing DMEM after 7 hours. The cells were scrapped out after 72 hrs, then centrifuged at 200g and cell pellet was subjected to lysis using 150 mM NaCl-20 mM Tris pH 8:0 buffer containing 10% sodium deoxy cholate. The lysate was then treated with Benzonase nuclease HC (Millipore) for 45 minutes at 37 ^o^C. Benzonase treated lysate was centrifuged at 3000xg for 15 mins and the clear supernatant was then subjected to HiTrap^®^ Heparin High Performance (GE) affinity column chromatography using a peristaltic pump. The elute from the Heparin column was then concentrated using Amicon Ultra centrifugal filters (Millipore). Titering of viral stock was determined as approximately 10^11^ infectious particles/mL.

### Injections of adeno-associated viruses

We injected AAVs carrying a cre dependent mCherry construct in one PV-IRES-Cre/R26REYFP intraparenchymally. The mouse was anaesthetized with isoflurane (4%, Nycomed, airflow 1 l/min) in an induction chamber. After induction, the mouse was moved to a stereotactic apparatus and placed on a heating pad (37°C) throughout the procedure. Eye ointment was applied to protect the cornea. The following analgesic was applied subcutaneously: buprenorphine hydrochloride (0.1 mg/Kg, Temgesic, Invidior), meloxicam (1 mg/Kg, Metacam Boerringer Ingelheim Vetmedica), bupivacaine hydrochloride (1 mg/Kg at the injection site, Marcain, Astra Zeneca). The head was fixed with ear bars before the surgery. The skin above the skull was shaved and disinfected with Pyrisept. An incision was made to expose the skull at the selected location. The skull was thinned with a drill and a small hole was made with a glass pipette at the appropriate coordinates to target the perirhinal cortex (from bregma: AP −3 mm, ML 4.4 mm, DV 1.3 and 2 mm). Pressure injections were performed with glass pipettes attached to an injector (Nanoliter 2010, World Precision Instruments) controlled by a microsyringe pump controller (Micro4 pump, World Precision Instruments). Two injections (200 nl each) along the dorsoventral axis were performed at 50 nl/minute. After retracting the pipette, the wound was rinsed with saline and sutured. The mouse could recover in a heated chamber before being returned to the home cage. The mouse received post operational analgesic 24 hours after the surgery (meloxicam 1 mg/Kg). The survival time for transduction and expression of the virus was 15 days.

### Histology

Mice were anaesthetized with isoflurane and euthanized with an injection of pentobarbital (i.p. 100 mg/Kg, Apotekerforeninger). The mice were subsequently perfused transcardially with Ringer’s solution (0.85% NaCl, 0.025% KCl, 0.02% NaHCO_3_) before perfusion with 4% paraformaldehyde (PFA) in PB (pH 7.4). After removing the brains from the skull, they were stored at 4°C in PFA for 2-3 hours and then moved in a solution containing sucrose 15% in PB and kept at 4°C overnight. The brains were moved to sucrose 30% in PB for two days before being sectioned at the freezing microtome. We cut sections of 50 μm, collected in six, equally spaced series in a solution containing: 30% glycerol, 30% ethylene glycol, 40% PBS. The series were stored at −20°C until used for immunostainings.

Slices were washed in PB (3 × 10 minutes) before being incubated in blocking solution (0.1% Triton-100X, 10% NGS in PB) for one hour. After blocking, the slices were moved to a new well and incubated for 2-3 days at 4°C in a solution with the primary antibodies (0.1% Triton-100X, 1% NGS in PB). After washing in PB (3 × 1 hour), the slices were incubated at 4°C overnight with the secondary antibodies. The slices were washed in PB (3 × 10 minutes) and then mounted on SuperFrost slides (Thermo Fisher Scientific) in PB, and left to dry overnight at RT. Once dried, the slides were washed in xylene before being coversliped with entellan in xylene (Merck Chemicals, Darmstadt, Germany). We used the following primary antibodies: Guinea Pig anti-NeuN (1:1000, Sigma Millipore, #ABN90P), Rabbit anti-parvalbumin (1:1000, Swant, #PV-27), Mouse anti-parvalbumin (1:1000, Sigma, #P3088), Rat anti-RFP (1:1000, Proteintech, #5f8), Chicken anti-GFP (1:1000, Abcam, #ab13970). We used the following secondary antibodies: Goat anti-guinea pig A647 (1:500, Invitrogen, #A-21450), Goat anti-Rat A-546 (1:500, Invitrogen, #A11081), Goat anti-Rabbit A488 (1:500, Invitrogen, #A11008), Goat anti-chicken A488 (1:500, Invitrogen, #A11039), Goat anti-rabbit A546 (1:500, Invitrogen, #A11010), Goat anti-rabbit A635 (1:500, Invitrogen, A31576).

### Image acquisition and analysis

We used a confocal microscope (Zeiss LSM 880 AxioImager Z2) to image regions of interest (ROIs) to count labeled neurons. Images of PER, wS1, PrL, IL and LEC were taken with a x20 air objective with 1 Airy unit pinhole size and saved as czi files. Files containing the ROIs were uploaded in Neurolucida (Micro Bright Field Bioscience) for analysis. We created contours to delineate the layers of the ROIs using the NeuN signal. The PER was delineated according to (Beaudin et al., 2013), LEC was delineated according to (Ohara et al., 2021), PrL and IL were delineated according to (Van De Werd et al., 2010). Markers for each signal were used to count cells labeled by different markers. Quantifications were done in Neurolucida explorer (Micro Bright Field Bioscience) and exported in excel files. The density of neurons labeled by a marker was measured as the number of labeled neurons in a contour divided by the area of the contour (cells/mm^2^). The specificity measures the percentage of cells labeled by the mouse line that co-express PV and was measured dividing the number of RFP-expressing neurons in Ai9 or GFP-expressing neurons in R26REYFP co-expressing PV by the number of RFP-expressing neurons or GFP-expressing in R26REYFP. The efficiency measures the percentage of PV-expressing neurons labeled by the mouse line and was calculated as the number of RFP-expressing neurons in Ai9 (or GFP-expressing in R26REYFP) co-expressing PV divided by the number of PV-expressing neurons. We quantified between 2 and 5 sections per cortical region. All graphs were created in excel, images of ROIs were created in Zen lite (Zeiss) and figures in illustrator.

Data for the comparison of the PV-IRES-Cre and PV-T2A-Cre were obtained from the Allen Institute website: Mouse Brain Connectivity Atlas, transgenic characterization. Samples for the PV-IRES-Cre: 100141217 Cre ISH, 81709692 td-Tomato ISH, 81657984 FISH. Samples for the PV-T2A-Cre: 100141203 Cre ISH, 81658019 td-Tomato ISH, 81811663 FISH.

## Results

We used immunofluorescence for PV on tissue from mice expressing tdTomato in PV-INs to address the specificity and efficacy of the PV-IRES-Cre mouse line in the mouse PER. The distribution of tdTomato across the cortex recapitulates the pattern of PV expression: i) higher expression in thalamo recipient layers (4 and 5B) in sensory cortex; ii) a bias towards deep layers throughout the cortex; iii) higher density in dorsal cortex than in ventral cortical areas (Fig. 1) (Tremblay et al., 2016).

**Figure 1.**
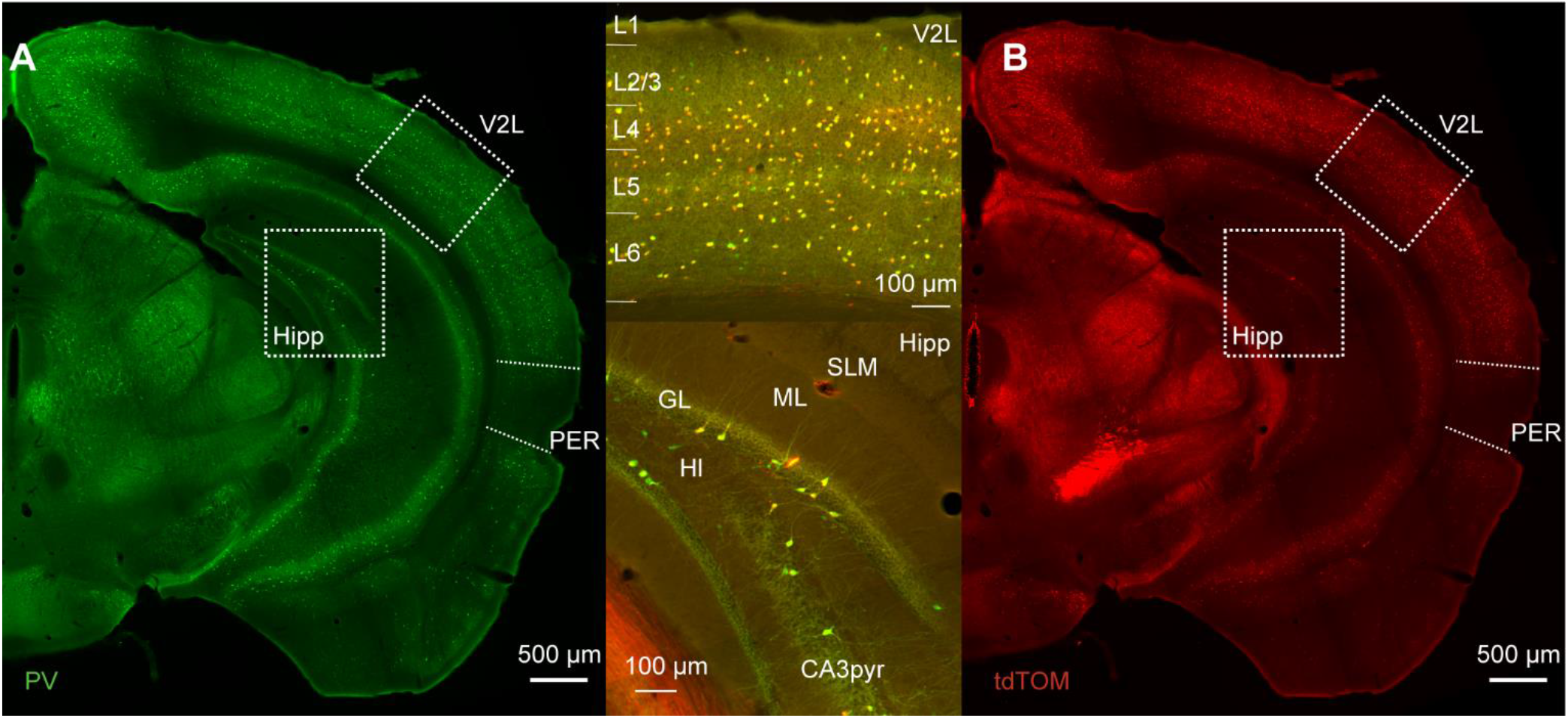
A. Representative immunofluorescence staining for PV of a slice at about bregma −3 mm of a PVcre/Ai9 mouse. Dotted boxes highlight the lateral part of the secondary visual cortex (V2L) and the hippocampus (Hipp). The perirhinal cortex (PER) is delineated by the dotted line on the lateral side of the brain. B. Representative image of the immunofluorescence for tdTomato in the same section as A. Upper insert shows the merged signals in the cortex, and lower insert shows merged signals in the hippocampus (SLM, stratum lacunosum-moleculare; ML, molecular layer; GL, granule layer of the dentate gyrus; HI, hilus; CA3pyr, pyramidal layer of the CA3 region).

We measured the density of PV-immunoreactive neurons (PV-IR) and tdTomato expressing neurons across the layers of PER. The pattern of tdTomato expression showed a bias towards the deep layers similarly to that of PV-IR neurons (Fig. 2A-B, D-E). However, the density of tdTomato expressing neurons was lower than that of PV-IR neurons (tdTomato: 122.56 ± 31.3 cells/mm^2^; PV-IR: 274.68 ± 10.9 cells/mm^2^, n= 4 mice) (Fig. 2A-B, D-E). Indeed, several PV-IR cells did not express tdTomato in the PV-IRES-Cre mouse line (Fig. 2C). The density of PV-IR neurons not expressing tdTomato was higher than that of tdTomato expressing neurons (Fig. 2F). This result suggests that the PV-IRES-Cre mouse line is very inefficient in labeling PV expressing GABAergic neurons in the mouse PER.

**Figure 2.**
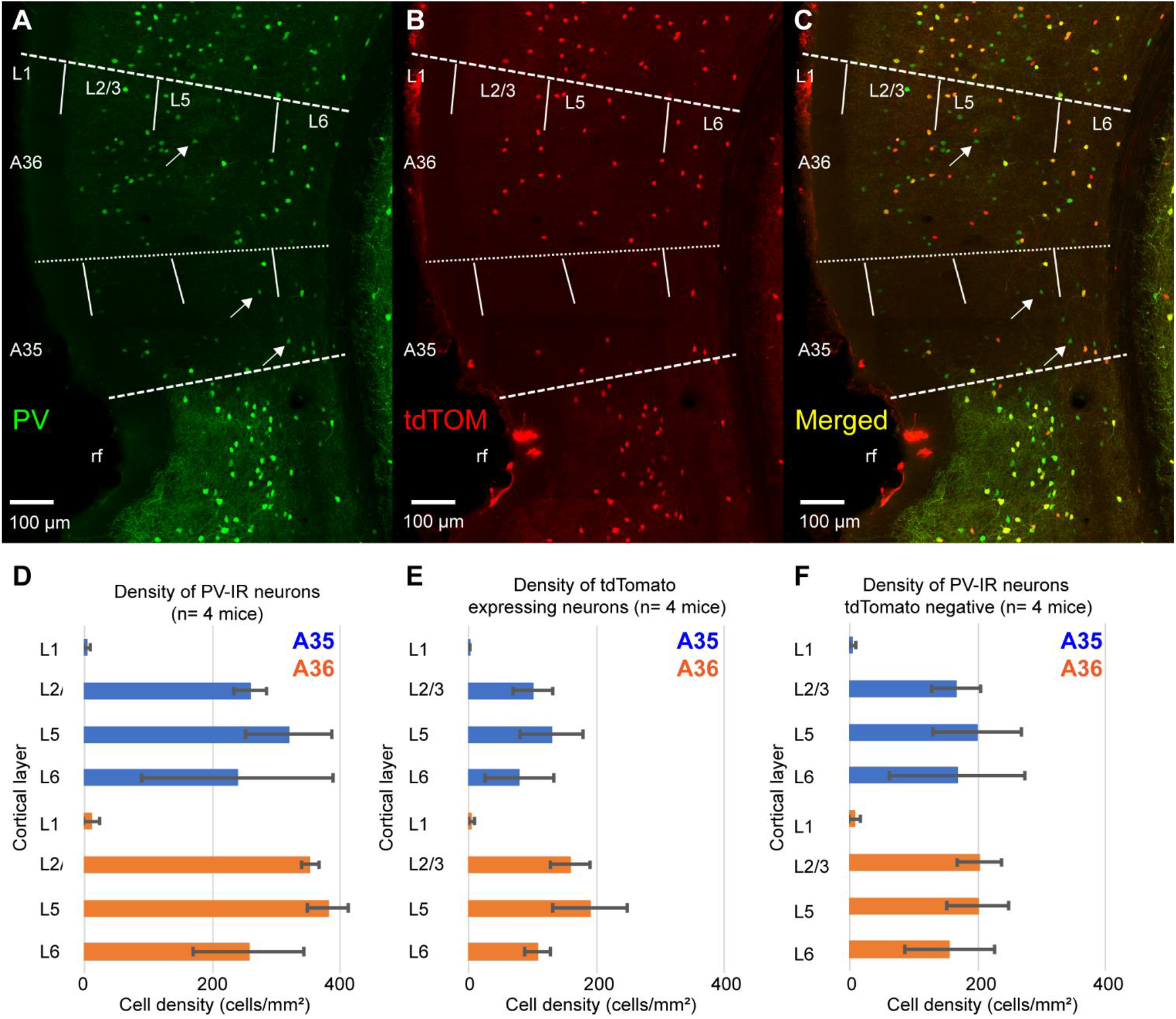
A-C. Representative confocal stacks showing immunofluorescence for PV (A), tdTomato (B) and the overlap of the two signals (C) in PER of a PV-IRES-Cre mouse. Arrows indicate PV-IR neurons that do not express tdTomato. rf, rhinal fissure. D. Bar graph showing the density of PV-IR neurons across the layers for A35 and A36. E. Bar graph showing the density of tdTomato-expressing neurons across the layers of A35 and A36. F. Bar graph showing the density of PV-IR that do not express tdTomato across the layers of A35 and A36.

We tested this hypothesis by quantifying the specificity (i.e., the percentage of tdTomato expressing neurons that also express PV) and the efficiency (i.e. the percent of PV-IR neurons labeled by tdTomato) of the PV-IRES-Cre mouse line in PER (Fig. 3). The majority of tdTomato expressing neurons was PV-IR, suggesting that the mouse line shows a high specificity for PV expressing neurons (Fig. 3A-B). However, we found that less than 50% of PV-IR neurons throughout the layers of PER were labeled by tdTomato (Fig 3A and C). These data demonstrate that the PV-IRES-Cre mouse line shows a low efficiency in labeling PV expressing neurons in PER. We confirmed our results using a different primary antibody against PV (mouse anti-PV) (Supp. Fig. 1). Moreover, the transgenic characterization of the Allen Institute for Brain Science demonstrates by fluorescent in situ hybridization (FISH) that many neurons expressing PV do not express tdTomato (Supp. Fig. 2).

**Figure 3.**
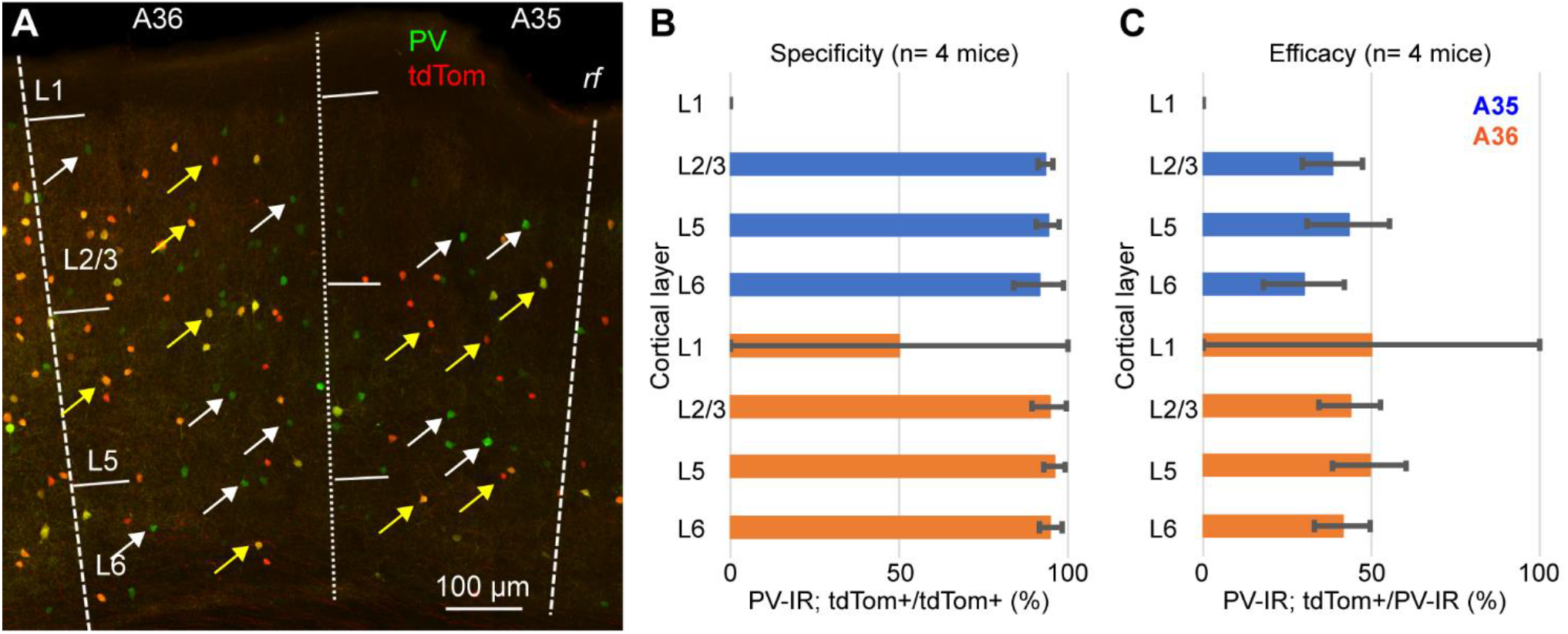
A. Representative confocal stack showing the overlap of immunofluorescence for PV (green) and tdTomato (red) in PER; yellow arrows indicate double labeled cells and white arrows indicate PV-IR cell that do not express tdTomato. B. Bar graph showing the specificity of the PV-IRES-Cre mouse line in PER in 4 animals. C. Bar graph showing the efficacy of the PV-IRES-Cre mouse line in PER in 4 animals.

As an internal control, we quantified the specificity and efficiency of the PV-IRES-Cre line in the barrel cortex. For these quantifications we used the same tissue as for the measurements in PER. As previously reported for the visual cortex (Atallah et al., 2012), the PV-IRES-Cre was very specific and efficient in labeling PV expressing neurons in the barrel cortex (Fig. 4). We found that 89.64 ± 7.5 % of tdTomato expressing neurons also expressed PV, and 93.97 ± 0.7 % of PV expressing neurons was labeled by tdTomato. These results suggest that our measurements are not a consequence of our histological procedures and are consistent with results obtained by FISH.

**Figure 4.**
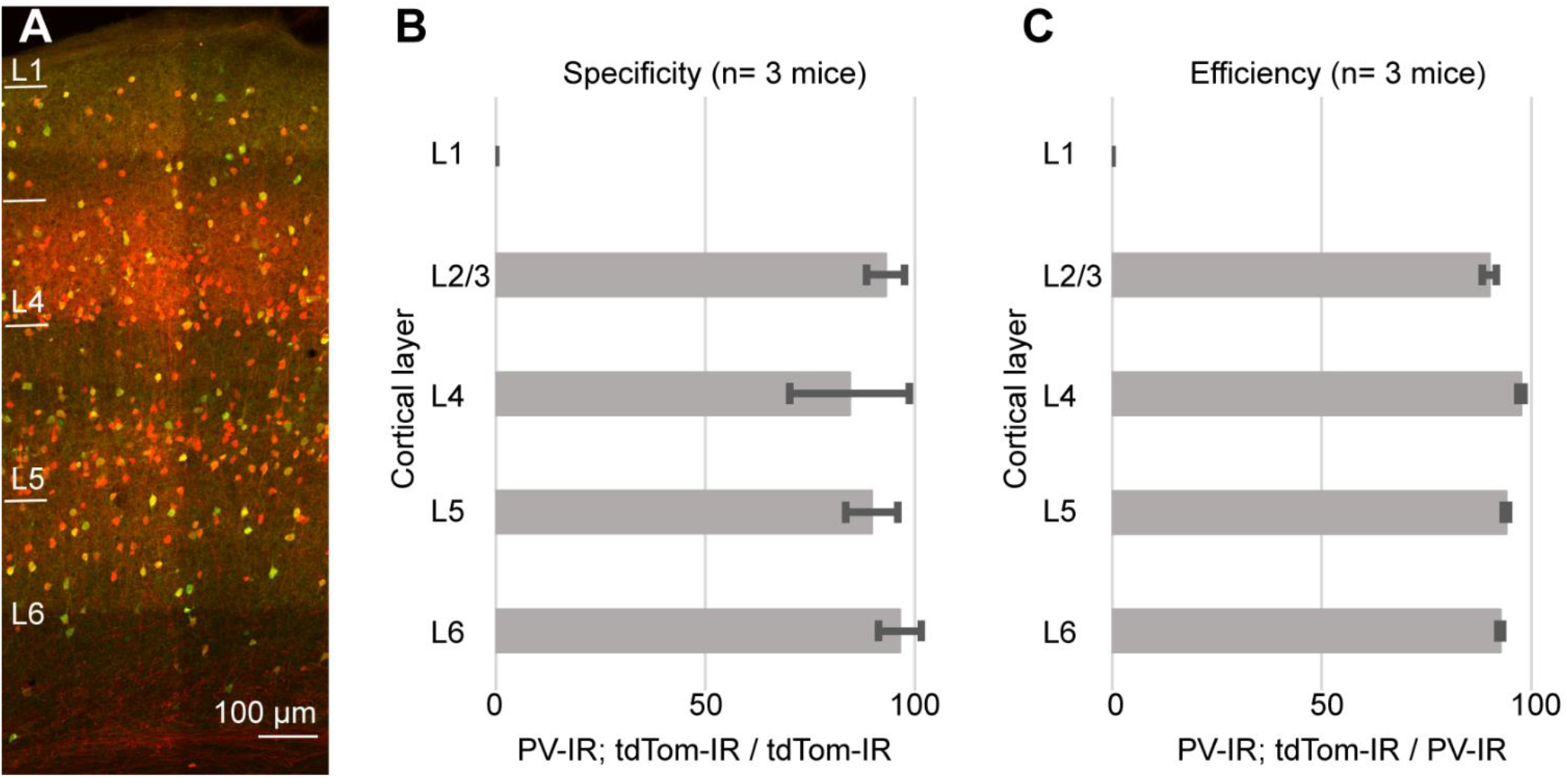
A. Representative confocal stack showing the overlap of immunofluorescence for PV (green) and tdTomato (Red) in barrel cortex. B. Bar graph showing the specificity of the PV-IRES-Cre mouse line in barrel cortex in 3 animals. C. Bar graph showing the efficiency of the PV-IRES-Cre mouse line in barrel cortex in 3 animals.

Our results can be explained either assuming that a population of PV-expressing interneurons does not express Cre-recombinase, or that the Ai9 reporter line is not reliable in PER. To test the reliability of the reporter line, we crossed the PV-IRES-Cre line with the Rosa26-stop-EYFP and measured the specificity and efficiency. Similar to our results with the Ai9 reporter line, the specificity was very high (96.85 ± 1 %, n= 2 mice) but only 49.15 ± 6.35 % of PV-IR neurons was labeled by YFP in PER (Fig. 5). Since both reporter lines exploit the Rosa26 locus, we hypothesized that this locus might be selectively silenced in a subpopulation of PV-expressing neurons in PER. We tested this hypothesis by injecting a Cre dependent virus AAV-FLEX-mCherry in PER, reasoning that if the hypothesis was correct then we would find PV-expressing neurons that also expressed mCherry but not YFP. However, we found that almost all mCherry expressing neurons also expressed YFP (94.79 %, n= 1 mouse, 96 mCherry cells), suggesting that the low efficiency resides in the PV-IRES-Cre mouse line rather than in the reporter line (Figure 6).

**Figure 5.**
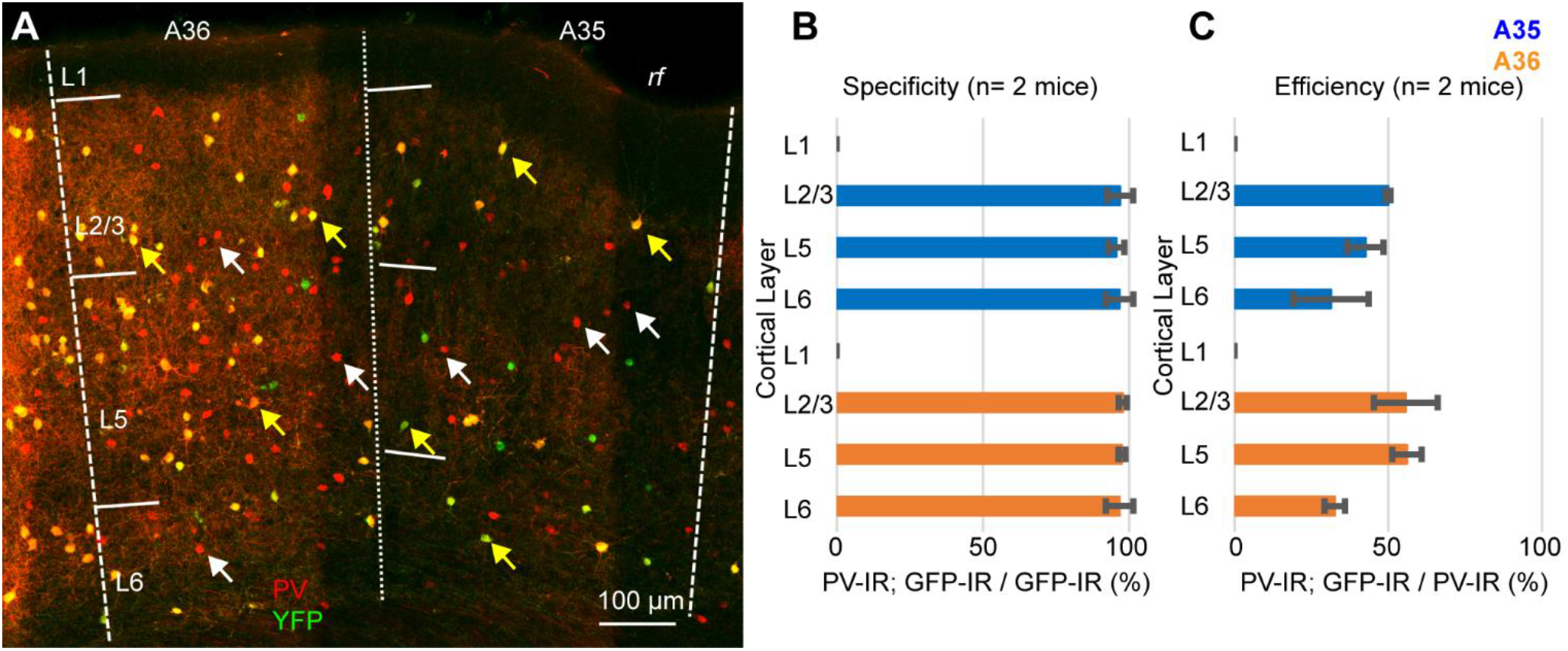
A. Representative confocal stack showing the overlap of immunofluorescence for PV (red) and YFP (green) in PER. Yellow arrows indicate double-IR neurons, white arrows indicate PV-IR neurons not labeled with YFP. B. Bar graph showing the specificity of the mouse line in 2 animals. C. Bar graph showing the efficiency of the mouse line in 2 animals.

**Figure 6.**
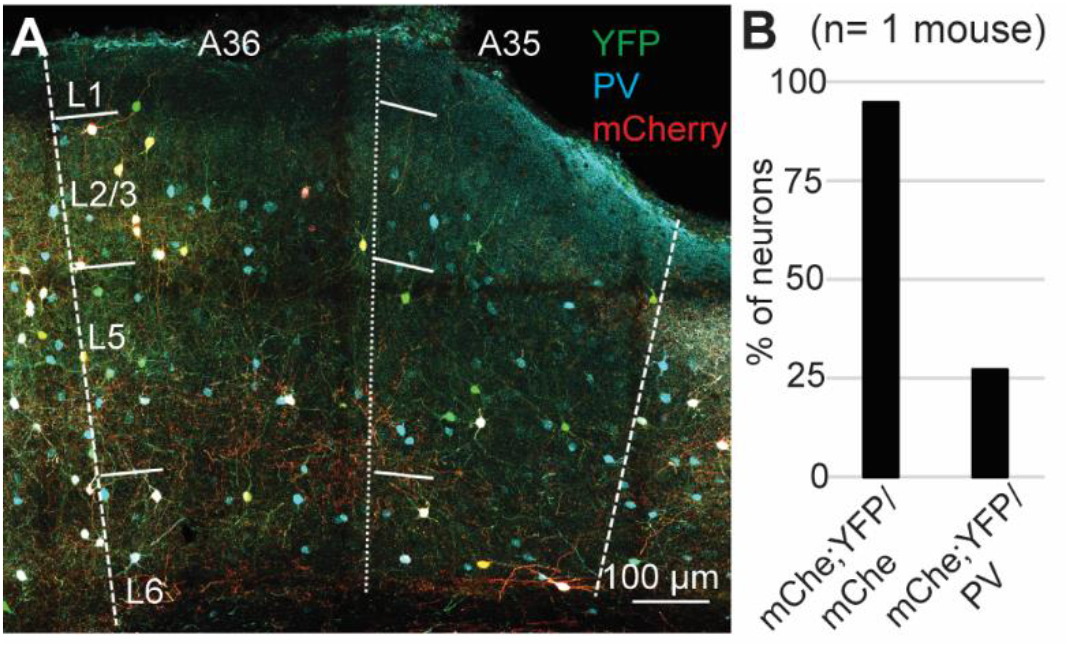
A. Representative confocal stack showing the overlap of PV (turquoise), YFP (green) and mCherry (red) in PER. B. Bar plot showing the percent of mCherry-expressing neurons that also expressed YFP, and the percent of PV-expressing neurons labeled by mCherry and YFP.

A recent report has demonstrated that the density of PV neurons is lower in associative areas of the cortex as compared to sensory-motor regions (Kim et al., 2017). These findings suggest that the low efficiency of the PV-IRES-Cre mouse line might be a shared feature of associative areas. To test this hypothesis, we measured the specificity and efficiency of the PV-IRES-Cre mouse line in three associative areas shown to have low density of PV neurons: prelimbic cortex (PrL), infralimbic cortex (IL) and LEC (Fig. 7). As reported for other cortical regions, the specificity of the PV-IRES-Cre mouse line was very high in all three areas (Fig. 7C-D), with the exception of L2 of IL where about 50% of tdTomato labeled neurons did not express PV (Fig. 7A1-3, C). The efficiency of the PV-IRES-Cre mouse line was lower than in the barrel cortex and similar to that reported in the PER (Fig. 3 and 7). These results suggest that the low efficiency of the PV-IRES-Cre mouse line is a phenomenon common to association cortices. The PV-IRES-Cre is the most widely used mouse line to label PV-INs, however another mouse line has been generated by the Allen Institute, the PV-T2A-Cre (Madisen et al., 2010). The PV-T2A-Cre line has been used to quantify PV-INs in the whole brain (Kim et al., 2017). These authors reported a lower density of PV-INs in PER as compared to sensorimotor areas. A qualitative comparison of the two mouse lines using available in situ hybridization data from the Allen Institute revealed low expression of Cre in both lines (Supp. Fig. 2). The expression of td-Tomato was higher in the PV-T2A-Cre than in the PV-IRES-Cre, and several td-Tomato cells do not seem to express PV (Supp. Fig. 2). The comparison of the two mouse lines suggests that Cre expression is low in both, and further examination of the expression of PV, Cre and td-Tomato is necessary in the PV-T2A-Cre.

**Figure 7.**
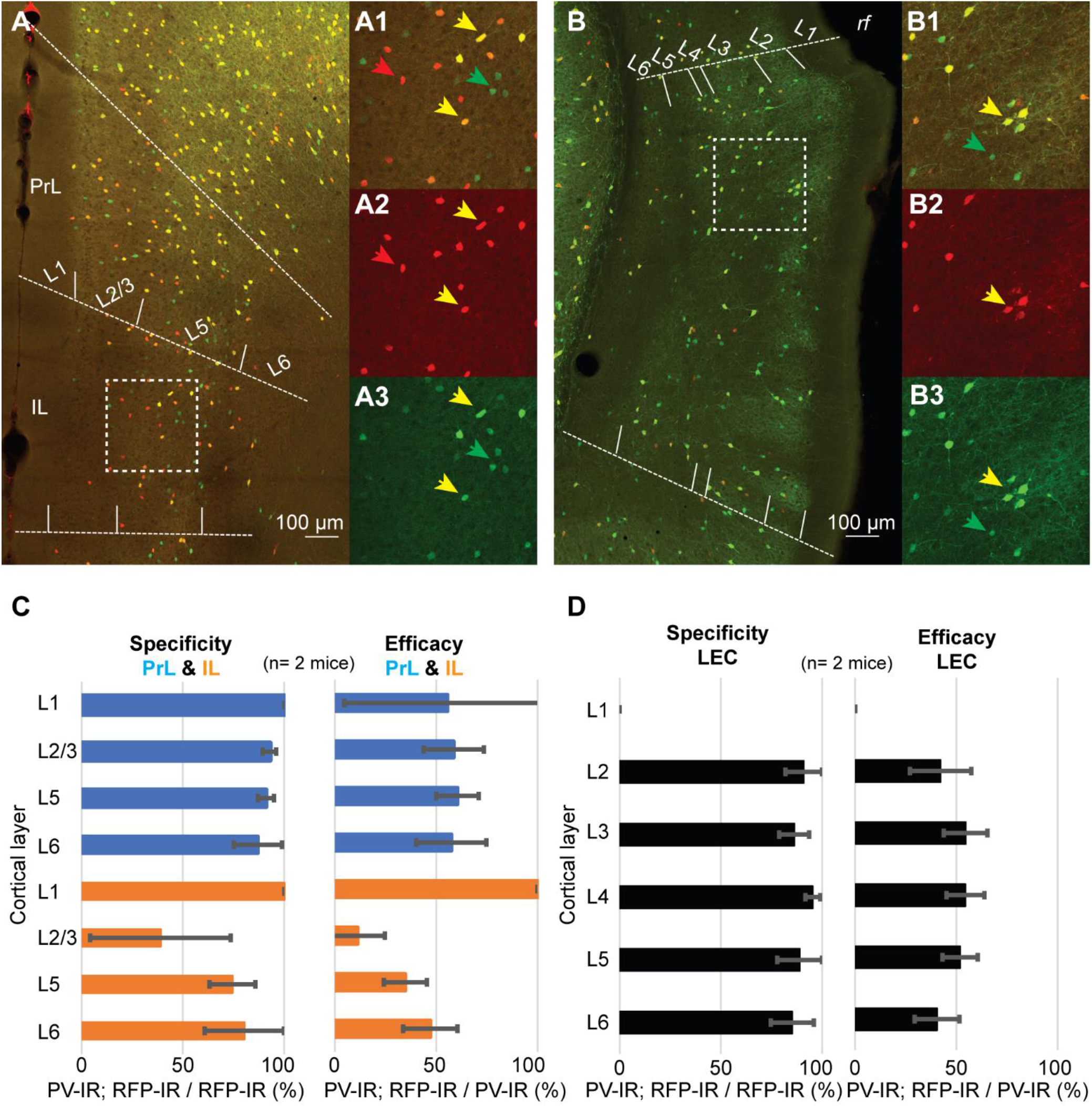
Representative confocal stack showing the overlap of PV (green) and tdTomato (red) in PrL and IL cortex (A), and in LEC (B). Inserts show magnification of the areas highlighted by dotted squares in A and B (A1-3, IL; B1-3, LEC). C. Bar graphs showing the specificity and efficiency of the PV-IRES-Cre mouse line in PrL and IL. D. Bar graphs showing the specificity and efficiency of the PV-IRES-Cre mouse line in LEC.

## Discussion

We demonstrate here that the PV-IRES-Cre mouse line has a low efficiency in labeling PV-INs in PER and this phenomenon might be a common feature of associative areas of the cerebral cortex. The PV-IRES-Cre mouse line has become an invaluable tool for studying fast-spiking basket cells across the telencephalon (Atallah et al., 2012; Yu et al., 2019; Straub et al., 2016). This mouse line has recently been used to estimate the number of PV-expressing fast-spiking basket cells in the cortex and hippocampus (Whissell et al., 2015). Our findings suggest that the density of PV-INs obtained by counting fate-mapped neurons in the PV-IRES-Cre mouse line is dramatically underestimated.

A recent study quantified labeled neurons in the PV-2A-Cre mouse line and showed that association areas of the cortex, including PER, contain a low density of PV-INs (Kim et al., 2017). The PV-IRES-Cre and the PV-T2A-Cre differ in regulatory elements, including the elements for multicistronic expression: IRES vs. 2A (Madisen et al., 2010). A qualitative examination of the characterization performed by the Allen Institute showed that the size of the labeled population is similar in the two mouse lines, suggesting that the low efficiency might be due to a difference within the population of PV-INs in PER. Interestingly, similar results to ours have been shown in the retina of PV-IRES-Cre mice (Gábriel et al., 2016). The retina of PV-IRES-Cre/YFP mice showed labeling in glial cells, but not in cell-types expressing PV (Gábriel et al., 2016).

PV-expressing fast-spiking neurons have also been implicated in controlling information flow along the cortico-hippocampal network. These neurons strongly respond to cortical inputs and provide feedforward inhibition to excitatory neurons in PER, controlling their output to LEC (Willems et al., 2018). Because of the high specificity of the PV-IRES-Cre line, we suggest that the PV-IRES-Cre mouse line can be used to study the cellular properties of fast-spiking basket cells in PER and their connectivity through paired recordings or circuits-mapping with soma-targeted optogenetics (Willems et al., 2018; Baker et al., 2016). However, the low efficiency of the PV-IRES-Cre mouse line might impair the optogenetic assessment of the circuits underlying the inhibitory control exerted by PER in the cortico-hippocampal network (e.g., using full-field light stimulation) (de Curtis and Paré, 2004).

Our work raises two questions: i) what are the mechanisms by which cells expressing PV do not express Cre?; ii) do labeled and unlabeled PV-INs belong to different subpopulations? The answer to the first question will require additional experiments to assess the activity of the locus containing Cre in labeled and unlabeled neurons in PER. The second question concerns with the biological implications of our findings in the context of neuronal diversity. Patch clamp recordings of PV-INs in the PER of PV-IRES-Cre mice show that these neurons express a fast-spiking phenotype (Willems et al., 2018). However, there is currently no information of the firing pattern of PV-INs not labeled by the mouse line. Assessing this lack of information requires a way to label either all GABAergic neurons or the entire PV-expressing population. One possibility is a comprehensive analysis of the firing patterns and molecular identity of GABAergic neurons in the GAD1-Cre mouse or in the GAD67-EGFP mouse. A transcriptomic analysis of the same mouse lines might further dissect subpopulations of the PV-INs in the PER.

The current observations do emphasize the importance of careful characterization of the tools used to study cell-types across cortical regions. Cortical regions might contain specialized populations not targeted by current transgenic or viral strategies or these strategies might show region specific bias in the labeled population.

---

**Supplementary figure 1.**
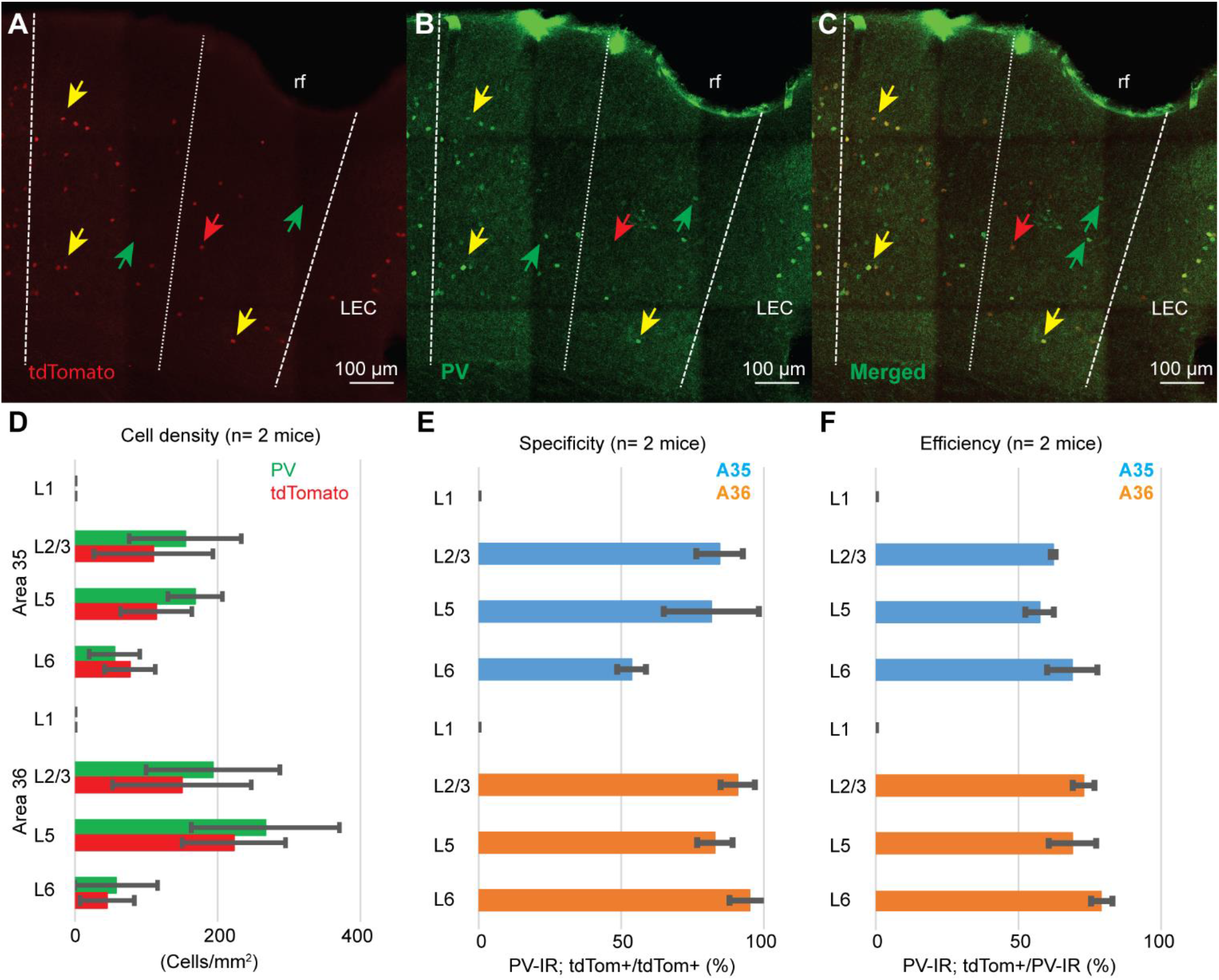
A-C. Representative immunostainings showing the expression of tdTomato (A, red), PV (B, green) and the merged image (C) in the perirhinal cortex of the PV-IRES-Cre line using a primary mouse anti-PV antibody. D. Bar plot showing the densities of PV immunoreactive (PV-IR) neurons (green) and tdTomato expressing neurons (red). E. Bar plot showing the specificity obtained with the mouse anti-PV. F. Efficiency obtained with the mouse anti-PV.

**Supplementary figure 2.**
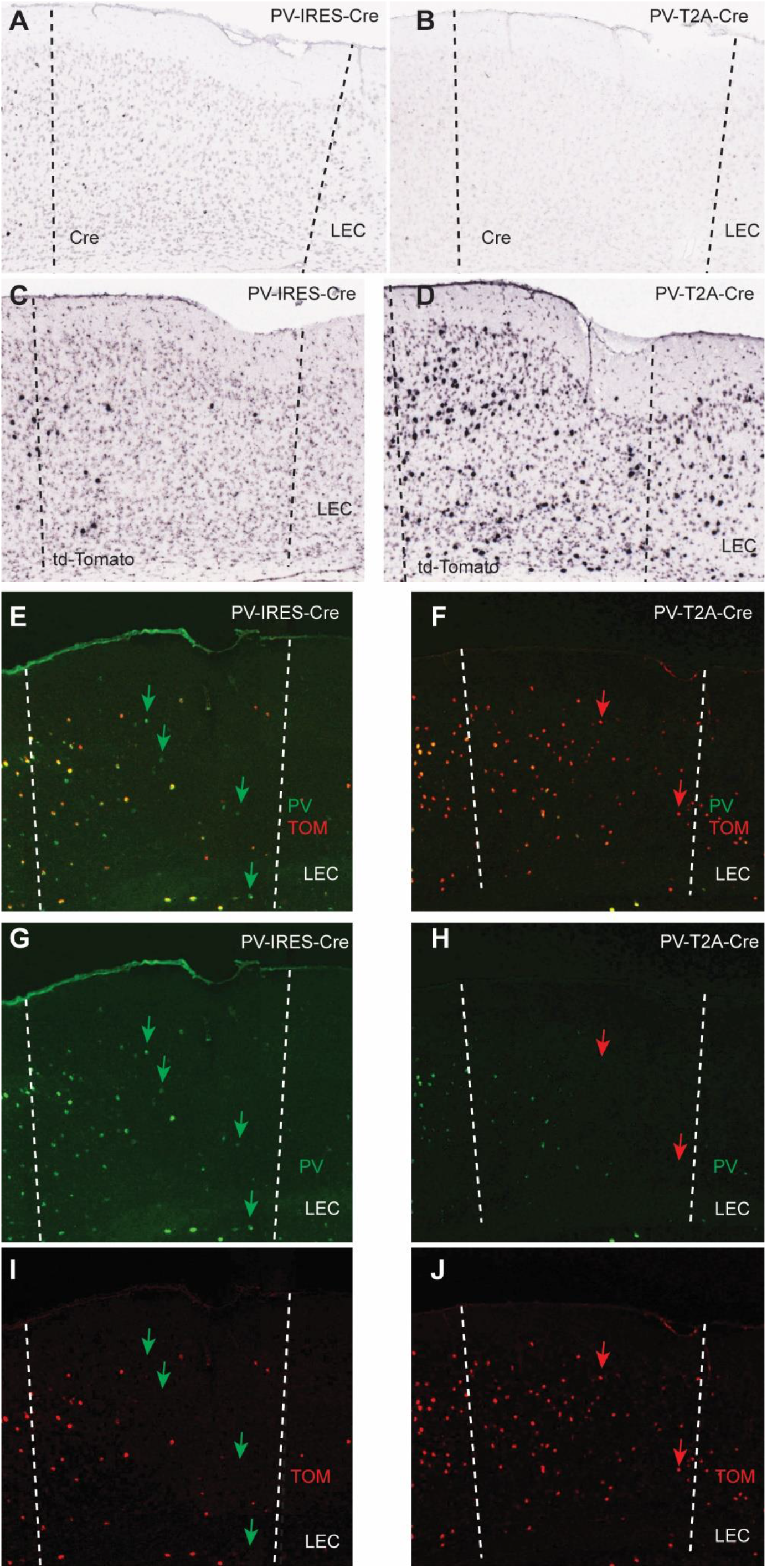
Comparison of the PV-IRES-Cre and PV-T2A-Cre mouse lines in the PER. A. PER of a PV-IRES-Cre mouse showing ISH for Cre. B. PER of a PV-T2A-Cre mouse showing ISH for Cre. C. PER of a PV-IRES-Cre mouse showing ISH for td-Tomato. D. PER of a PV-T2A-Cre mouse showing ISH for tdTomato. E-J. FISH for PV and td-Tomato in PV-IRES-Cre (E, merged; G, PV; I, td-Tomato), and in PV-T2A-Cre (F, merged; H, PV; J, td-Tomato). Green arrows point to PV-expressing neurons that do not express tdTomato. Red arrows point to td-Tomato neurons that do not express PV. LEC, lateral entorhinal cortex

